# Artificial neural network modelling of the neural population code underlying mathematical operations

**DOI:** 10.1101/2022.06.06.494909

**Authors:** Tomoya Nakai, Shinji Nishimoto

**Affiliations:** Center for Information and Neural Networks, National Institute of Information and Communications Technology, Suita, Japan; Lyon Neuroscience Research Center (CRNL), INSERM U1028 - CNRS UMR5292, University of Lyon, Lyon, France; Graduate School of Frontier Biosciences, Osaka University, Suita, Japan; Graduate School of Medicine, Osaka University, Suita, Japan

**Keywords:** fMRI, mathematics, IPS, encoding model, artificial neural network

## Abstract

Mathematical operations have long been regarded as a sparse, symbolic process in neuroimaging studies. In contrast, advances in artificial neural networks (ANN) have enabled extracting distributed representations of mathematical operations. Recent neuroimaging studies have compared distributed representations of the visual, auditory and language domains in ANNs and biological neural networks (BNNs). However, such a relationship has not yet been examined in mathematics. Here we used the fMRI data of a series of mathematical problems with nine different combinations of operators to construct voxel-wise encoding models using both sparse operator and latent ANN features. Representational similarity analysis demonstrated shared representations between ANN and BNN, an effect particularly evident in the intraparietal sulcus. Feature-brain similarity analysis served to reconstruct a sparse representation of mathematical operations based on distributed ANN features. Such reconstruction was more efficient when using features from deeper ANN layers. Moreover, latent ANN features allowed the decoding of novel operators not used during model training from brain activity. The current study provides novel insights into the neural code underlying mathematical thought.

## Introduction

Mathematical operations have long been regarded as a sparse, symbolic process in human neuroimaging studies. For example, neuroimaging data under subtraction and multiplication conditions was analysed based on sparse event-related designs including two operators (Ischebeck et al. 2006; Prado et al. 2011). In contrast, recent advances in artificial neural networks (ANN) enable extracting distributed representations of quantity information (Kim et al. 2021; Nasr, Viswanathan, and Nieder 2019) and mathematical operations (Schlag et al. 2019; Russin et al. 2021), where each operator is represented as a vector in a high-dimensional latent feature space. In particular, a recent study demonstrated that the intermediate layers of the tensor-product (TP)-transformer model can capture the compositional structure of mathematical expressions (Russin et al. 2021).

An emerging field of cognitive computational neuroscience has reported evidence on whether such latent features can explain the symbolic representations in the human brain. Previous studies has demonstrated that ANNs and biological neural networks (BNNs) partially share representations for visual (Yamins et al. 2014; Horikawa and Kamitani 2017; Kietzmann et al. 2019; Güçlü and van Gerven 2015; Groen et al. 2018), auditory (Kell et al. 2018; Koumura, Terashima, and Furukawa 2019) and language processing (Schrimpf et al. 2021; Hannagan et al. 2021; Goldstein et al. 2022; Caucheteux and King 2022; Schmitt et al. 2021). However, to date, the representational relationship between ANN and BNN in mathematics has not been investigated.

Previous studies used the encoding model approach (Naselaris et al. 2011) to compare the representational relationship between ANNs and BNNs (Schmitt et al. 2021; Ratan Murty et al. 2021; Jain and Huth 2018; Caucheteux and King 2022; Schrimpf et al. 2021; Goldstein et al. 2022). Encoding models quantitatively predict brain activity based on a combination of features extracted from the presented stimuli. Researchers have adopted this approach to comprehensively examine visual (Kay, Naselaris, et al. 2008; Nishimoto et al. 2011; Naselaris et al. 2009), auditory (Norman-Haignere, Kanwisher, and McDermott 2015; Nakai, Koide-Majima, and Nishimoto 2021), semantic (Huth et al. 2012, 2016; Popham et al. 2021), emotional (Koide-Majima, Nakai, and Nishimoto 2020; Horikawa et al. 2020) and many complex cognitive functions (Nakai and Nishimoto 2020). Encoding models also allow the comparison between sparse and latent features in the same brain data using feature-brain similarity (FBS) (Nakai, Koide-Majima, and Nishimoto 2021).

To investigate the relationship between the symbolic and distributed representations underlying mathematical operations in the brain, we used part of a 3-h fMRI dataset where subjects performed a series of mathematical problems with nine different combinations of operators (**Figure 1A**). By using sparse operator features, we constructed a voxel-wise encoding model to predict the brain activity specific to each mathematical operation (**Figure 1B**). We then extracted the latent features from the intermediate layers of an ANN (TP-Transformer) (Schlag et al. 2019) and constructed another encoding model based on these latent features. The prediction performance of each encoding model was evaluated using the remaining dataset. Next, we compared the representational relationship between the brain and ANN using representational similarity analysis (RSA). A brain representation of the sparse operator model was reconstructed by FBS analysis (Nakai, Koide-Majima, and Nishimoto 2021). Furthermore, these latent ANN features enabled decoding mathematical problems from brain activity, even for novel operators not included during model training. The current study provides new insights into the neural population code of mathematical thought.

**Figure 1.**
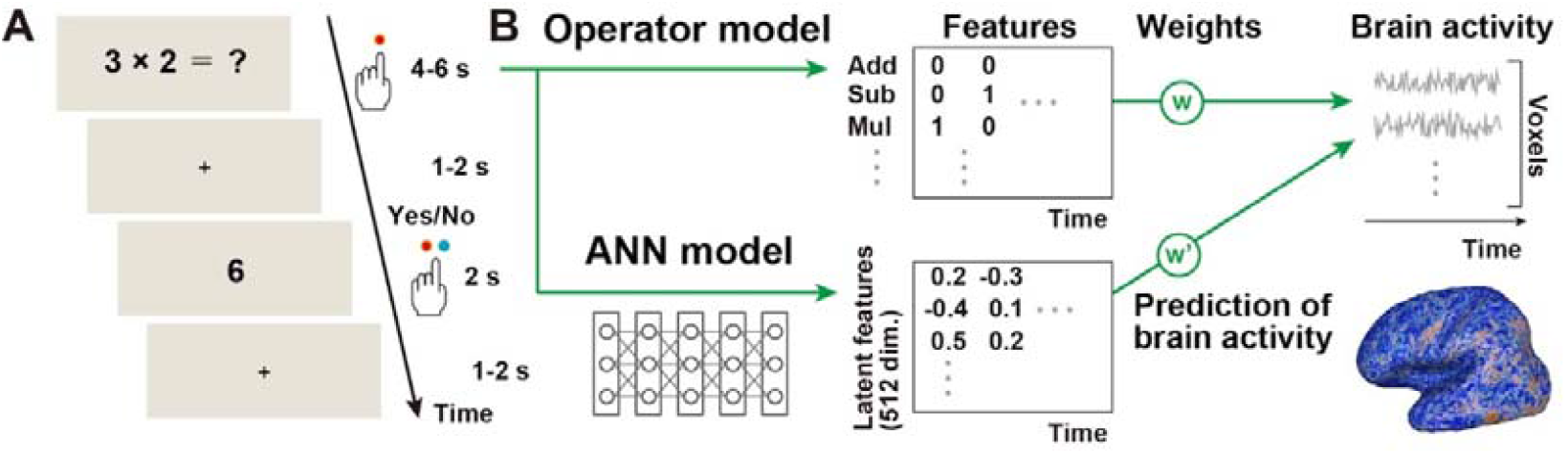
Experimental design. (**A**) Mathematical expressions were presented for 4 s for single-operator conditions and 6 s for double-operator conditions while the evoked brain activity was measured using fMRI. Subjects pressed a button on the left when they solved the given problems. The subjects again pressed the left or right button to indicate whether the probe digit stimulus (presented for 2 s) matched their answer. (**B**) Sparse operator and continuous artificial neural network features were extracted from mathematical problems. For each type of features, a voxel-wise encoding model was constructed to predict brain activity. Model weights were estimated using L2-regularised linear regression.

## Results

### Latent features extracted from ANN predicted the brain activity induced by mathematical problems

To investigate whether the categorical operator and latent ANN features capture brain representations of math problem solving, we constructed voxel-wise encoding models with both types of features using the training dataset and quantified the prediction accuracy of each model with the test dataset. Both models significantly predicted the activity of large brain regions of the bilateral frontal, parietal and occipital cortices (operator model, prediction accuracy across whole cortex = 0.040 ± 0.026 (mean ± s.d.), 21.5 ± 9.4% of voxels were significant; ANN model, prediction accuracy = 0.045 ± 0.025, 22.8 ± 10.0% of voxels were significant; two-tailed Wilcoxon signed-rank test across subjects, *p* = 0.078; **Figure 2A, S1**). The overall prediction maps between operator and ANN models were similar across all cortical voxels (Spearman’s correlation coefficient, *ρ* = 0.783 ± 0.088; **Figure 2B, S2**).

**Figure 2.**
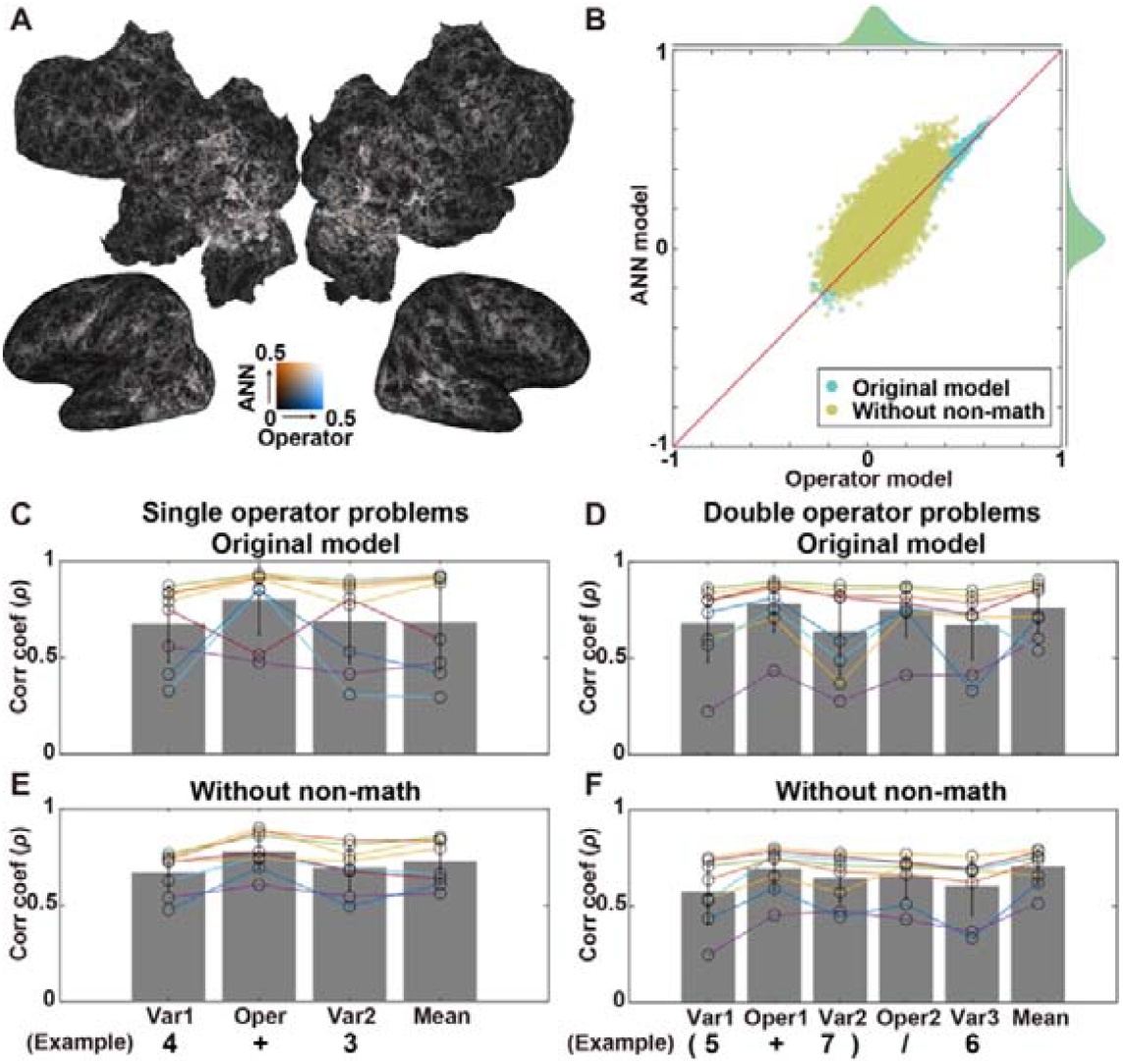
Prediction of brain activity by the ANN and operator encoding models. **(A)** Prediction accuracies of the sparse operator (blue) and latent ANN (red) models shown on the cortical surface of subject ID01. Only voxels with significant prediction are shown *(p* < 0.05, FDR corrected). (**B**) Scatter plot of prediction accuracies of ANN and operator models, shown for both the original model (cyan) and after exclusion of non-mathematical regressors (yellow). (**C-F**) Correlation between the prediction map of the operator model and those of ANN models, constructed using features from different positions, shown for both single- and double-operator problems in the original model (**C, D**) and after exclusion of non-mathematical regressors (**E, F**). Individual subjects’ data are indicated by small circles. Error bar, SD.

The above results suggest that both operator and ANN encoding models equivalently predict brain activity during math problem solving. However, these models may partially contain non-mathematical information. To exclude the influence of non-interesting features, we constructed additional encoding models concatenating the target features with other non-mathematical features such as motion energy (visual), reaction times (procedural load), number of letters (orthographic), word2vec (semantic) and button responses (motor). The ANN model outperformed the categorical model when those non-mathematical features were regressed out (operator model, prediction accuracy = 0.028 ± 0.024, 18.3 ± 8.0% of voxels were significant; ANN model, prediction accuracy = 0.043 ± 0.026, 21.7 ± 10.4% of voxels were significant; Wilcoxon signed-rank test, *p* = 0.039; **Figure 2B, S2**). Once again, the prediction maps between operator and ANN models were positively correlated (*ρ* = 0.667 ± 0.078), indicating that the ANN model captures latent information specific to mathematical operations.

Next, we examined how the information contained in the ANN differs depending on math expression components. We constructed ANN models using features from different positions in math expressions (e.g. ‘4’ and ‘+’ from ‘4 + 3’) and calculated the Spearman’s correlation coefficients between the prediction accuracy maps of the operator model and various ANN models. We found larger correlation coefficients for most subjects when using ANN features from operators for both single- and double-operator problems (Difference of Spearman’s correlation coefficients, single-operator problems, 0.121 ± 0.244, two-tailed Wilcoxon signed-rank test across subjects, *p* = 0.148; double-operator problems, 0.102 ± 0.070, *p* = 0.008; Figure **2C**, **2D**). This effect was particularly evident when non-mathematical features were regressed out (Difference of Spearman’s correlation coefficients, single-operator problems, 0.096 ± 0.059, two-tailed Wilcoxon signed-rank test across subjects, *p* = 0.008; double-operator problems, 0.068 ± 0.042, *p* = 0.008; **Figure 2E, 2F**). These results demonstrate that the ANN model captures representations of different math operators.

### The ANN model captured categorical representations of different math operations

To interpret the latent features of the ANN model and compare it with the sparse operator model, we performed an RSA (**Figure 3A**). In each anatomical region of interest (ROI), we calculated a representational similarity matrix (RSM) with the correlation distance among nine operators using the operator model (brain-based RSM) and another using ANN features (feature-based RSM). We found a significant correlation between the upper triangular matrices of brain-based and ANN feature-based RSMs in the bilateral frontal, parietal, temporal and occipital cortices (**Figure 3B**). Among cortical ROIs, we observed the largest correlation coefficients in the bilateral intraparietal sulci (IPS; left IPS, *ρ*, = 0.418 ± 0.093; right IPS, *ρ* = 0.416 ± 0.066).

**Figure 3.**
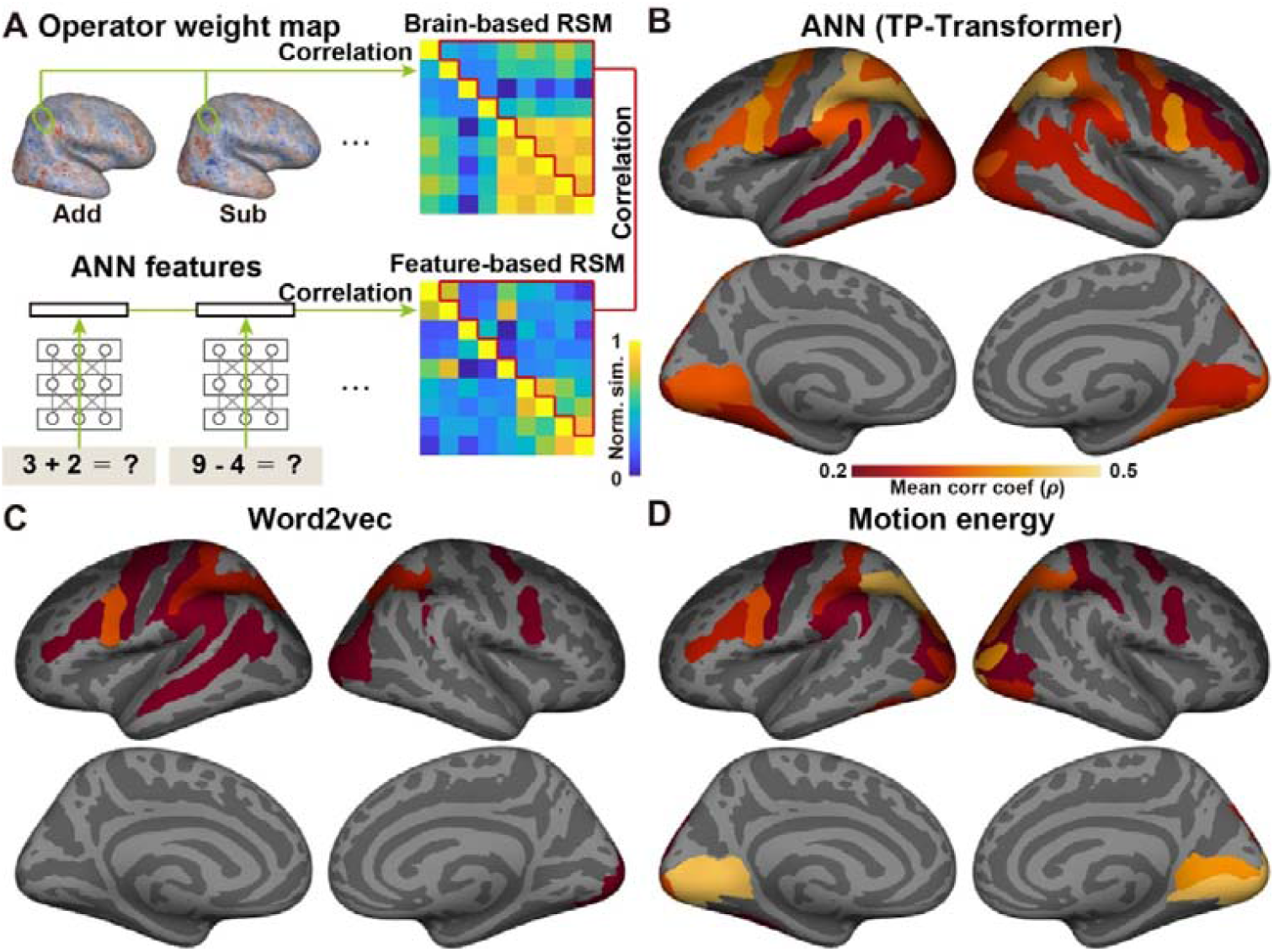
Similar brain representations in the sparse operator and latent ANN feature models. (**A**) Schematic description of representational similarity analysis. For both operator model weights and ANN feature vectors, the representational similarity was calculated including all condition pairs. The resultant brain-based and feature-based representational similarity matrices were further compared using Spearman’s correlation. (**B-D**) The Spearman’s correlation coefficient was calculated in each anatomical region of interest using (**B**) TP-Transformer, (**C**) word2vec and (**D**) motion-energy features.

As control analyses with non-mathematical latent features, we also calculated feature-based RSMs using other latent features of the word2vec (semantic) and motion-energy (visual) models. The word2vec is a widely-known model of semantic information used in previous studies in encoding/decoding language models (Nishida and Nishimoto 2018; Nishida et al. 2021). The motion-energy model is used to reconstruct visual information from brain activity (Nishimoto et al. 2011). The word2vec model showed a left-lateralised pattern and larger correlation coefficients in the left inferior frontal and superior temporal cortices (**Figure 3C**), whereas the motion-energy model showed the largest correlation coefficients in the bilateral occipital cortex (**Figure 3D**). These results suggest that different types of latent features may capture different representational aspects of mathematical operations.

### Latent ANN features could reconstruct brain representations of the sparse operator model

Although RSA calculates similarity in an abstract space, it does not directly assess whether there is a relationship between latent features and brain representations. Moreover, RSA calculates similarity based on geometric multivoxel patterns and does not provide similarity information for each voxel. To address these issues, we performed an FBS analysis as previously described (Nakai, Koide-Majima, and Nishimoto 2021). Specifically, we calculated the Pearson’s correlation coefficient between the reference vector of each operator and the weight vector extracted in each cortical voxel (**Figure 4A**).

**Figure 4.**
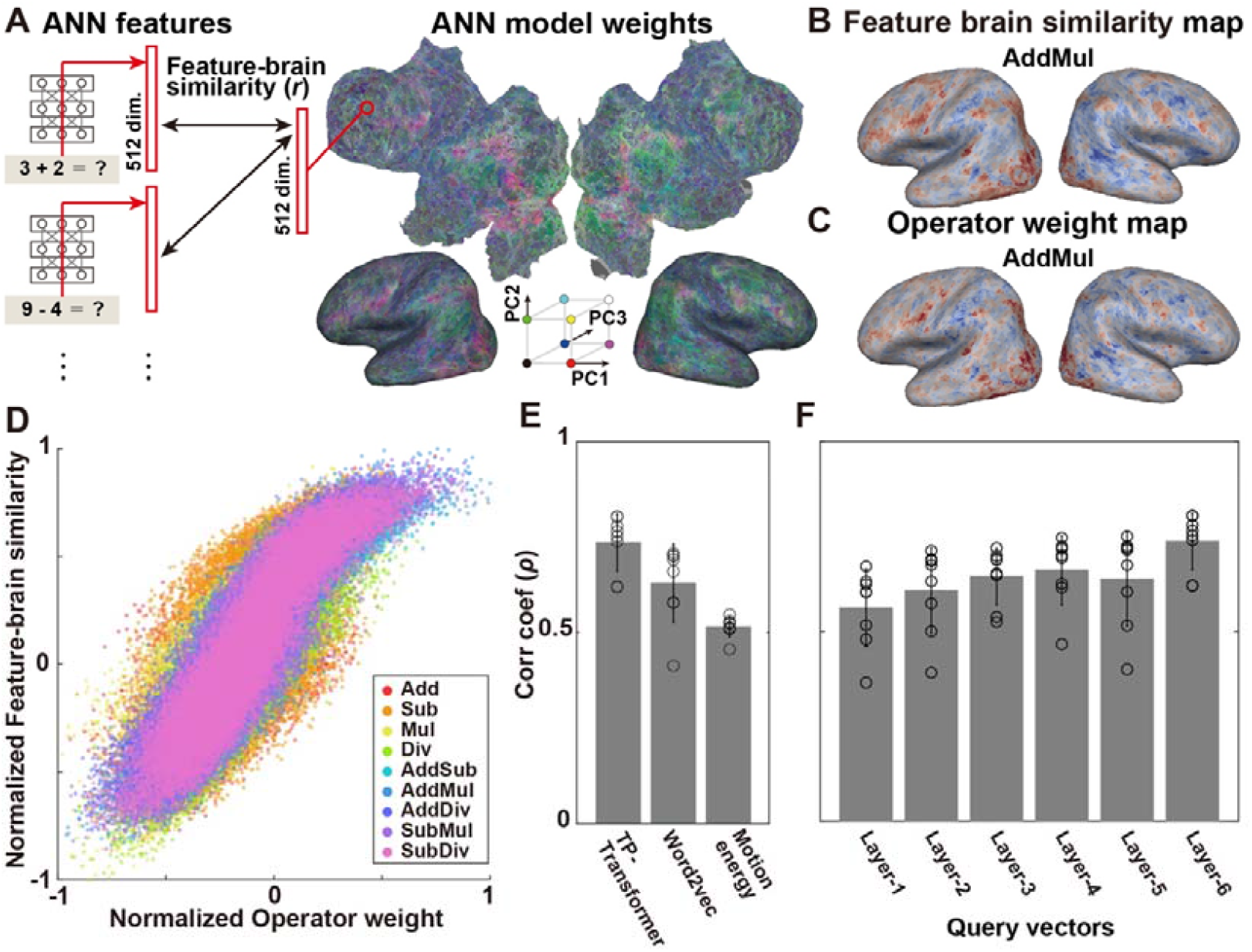
Reconstruction of the sparse operator model space based on latent ANN features. (**A**) Schematic illustration of feature-brain similarity (FBS) analysis. The ANN model weight map is visualised with three top components obtained using principal component analysis in red, green and blue. (**B**) FBS map obtained for the AddMul condition, shown for subject ID01. (**C**) Weight map of the AddMul condition obtained using the sparse operator model. (**D**) Scatter plot of all cortical voxels showing a correlation between operator weight and FBS values for each condition. (**E**) Comparison of different latent feature models. Correlation coefficients between operator weights and FBS values calculated using whole-cortical voxels plotted for the ANN (TP-Transformer), word2vec and motion-energy models. (**F**) Comparison of correlation coefficients between operator weights and FBS values obtained using query vectors from different ANN encoding layers (layers 1–6).

This way, we obtained a single FBS value for each cortical voxel and for each operator. The resultant FBS map obtained using latent ANN features (**Figure 4B**) was similar to the weight map of the sparse operator model (**Figure 4C**). We found a significant correlation between the FBS map and the operator-weight map (mean correlation coefficient across nine operators, 0.724 ± 0.031, *p* < 0.001: **Figure 4D, S3**). By comparing different latent features, we found that ANN-extracted features (TP-transformer) outperformed those extracted from motion energy (visual) and word2vec (semantic) models using whole-cortical voxels (Wilcoxon signed-rank test; *p* = 0.008) (**Figure 4E**). Furthermore, the correlation between the operator-weight map and FBS values increased with ANN hierarchy (difference between encoding layer-6 and other layers, Wilcoxon signed-rank test, *p* < 0.008; **Figure 4F, S4**). These results indicate that the latent ANN features can serve to reconstruct the brain representations of math operators obtained with categorical features.

### The ANN model enables decoding novel math problems

To further examine the specificity of the ANN model, we performed a series of decoding analyses based on ANN features (**Figure 5A**). First, the generalizability of decoding models was evaluated by training a decoding model using eight of the nine operators in the training dataset (e.g. ‘Add’, ‘Sub’, ‘Mul’, ‘Div’, ‘AddMul’, ‘AddDiv’, ‘SubMul’ and ‘SubDiv’) and testing it with the remaining operator in the test dataset (e.g. ‘AddSub’). Second, decoding accuracy was calculated as the area under the receiver operating characteristic (ROC) curve (AUC, **Figure S6**) (Nishida and Nishimoto 2018). As a result, mathematical problems were significantly decoded for all subjects (mean decoding accuracy across nine operators, 0.724 ± 0.031; one-sided permutation test for average decoding accuracy, *p* < 0.001, FDR corrected; **Figure 5B**).

**Figure 5.**
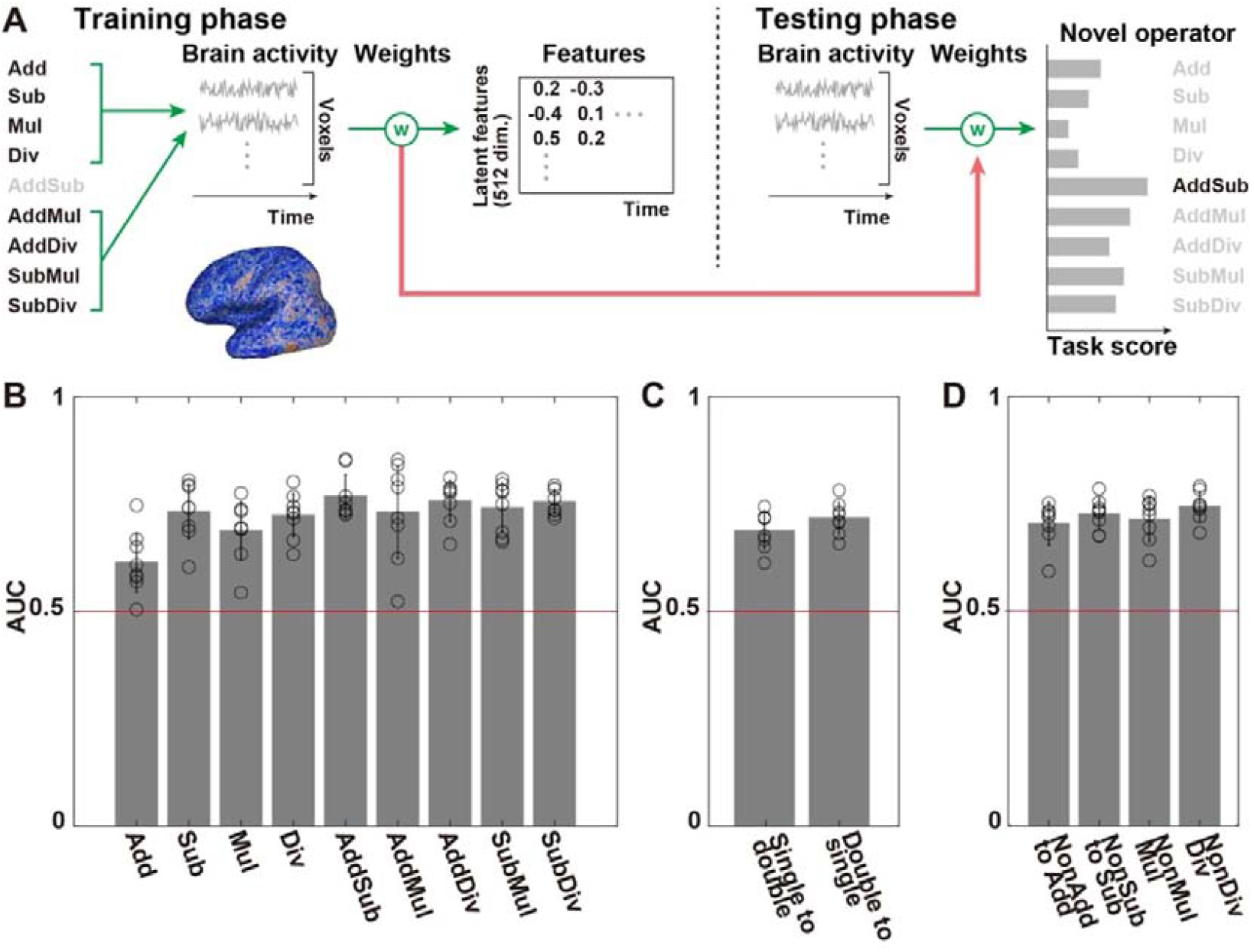
Decoding math problems from brain activity using ANN features. (**A**) The decoding model was constructed using eight out of nine operator problems in the training dataset and tested with the remaining (novel) operator in the test dataset. Decoding accuracy was evaluated by comparing task scores. (**B**) Decoding results using novel operators. Decoding accuracy evaluated using the area under the receiver operating characteristic curve. Receiver operating characteristic curves for individual subjects are shown in Supplementary information (**Figure S5**). Chance level accuracy (0.5) is indicated by a red line. Individual subjects’ data are indicated by small circles. Error bar, SD. (**C**) Decoding models trained using single-operator problems and tested with double-operator problems (‘single to double’) and vice-versa (‘double to single’). (**D**) Decoding models trained using problems without target operators and tested using problems with the target operator. Four different models were tested (‘NonAdd to Add’, ‘NonSub to Sub’, ‘NonMul to Mul’ and ‘NonDiv to Div’) depending on the four target operators (‘Add’, ‘Sub’, ‘Mul’ and ‘Div’), respectively.

Next, we investigated if the generalizability of decoding accuracy was confined to a specific problem structure (i.e. single- or double-operators), as math problem structure may affect brain activity patterns (Nakai and Sakai 2014). We thus trained decoders using single operators in the training dataset and tested with double-operators in the test dataset; inversely, decoders trained using double-operator problems were applied to single-operator problems in the test dataset. In both cases, the decoding accuracy was larger than chance (0.5) for all subjects (mean decoding accuracy across nine operators, 0.670 ± 0.044; *p* < 0.001, FDR corrected; **Figure 5C**).

Although the above analysis significantly decoded operators not used for model training, since part of the single operators in the training dataset (‘Add’ and ‘Mul’) were also used among the double-operators in the test dataset (e.g. ‘AddMul’), we constructed additional decoding models without a target operator (e.g. without ‘Add’, ‘AddSub’, ‘AddMul’ and ‘AddDiv’) and tested them with the target operator (e.g. ‘Add’). We again found significant decoding accuracy for all four target operators (mean decoding accuracy across nine operators, 0.721 ± 0.034; *p* < 0.001, FDR corrected; **Figure 5D**). These results indicate that the ANN model captures detailed representations of different mathematical problems.

## Discussion

In the current study, we constructed voxel-wise encoding models using both a sparse operator and latent ANN features for math problem solving and examined the representational relationship in the brain between these two features types. Although the two models showed similar prediction maps, the ANN model was more robust than the operator model after controlling for various non-mathematical features. Representational similarity and FBS analyses demonstrated shared representational patterns between ANN and BNN. This effect was particularly evident in the IPS. Latent ANN features further allowed decoding of novel operators not used for model training. These results indicate that ANN-based distributed representations in the brain partially explain the symbolic processing of mathematical operations.

Although we did not use any control condition, we excluded the possible effects of non-mathematical features (sensorimotor, language, executive load) by concatenating motion energy (visual), reaction times (procedural load), number of letters (orthographic), word2vec (semantic) and button responses (motor) features to the original feature matrix (**Figure 2B**). The outperformance of the ANN model in this control analysis may imply that the latent ANN model captures more detailed information on math problem solving across the cortex. Conversely, the sparse operator model might be a coarse representation of such complex cognitive processes, insufficient to explain the brain activation patterns of mathematical operations. The control analysis also indicates that the human brain has a specific population coding of mathematical operations independent of sensorimotor, language, or general executive load.

Correlation analyses of prediction maps demonstrated similar prediction patterns between the sparse operator and latent ANN models based on math operators (**Figure 2C, D**). These results are consistent with a previous study using the same ANN model (Russin et al. 2021), which showed that the feature vectors of intermediate calculations of a math expression were more similar to feature vectors extracted from operators of the same expression than those from digits. Our results are also consistent with the idea of sparse operator features, which assumes different representations across operators, regardless of the digits used.

The RSA revealed different similarity patterns between the sparse operator model and three types of latent features (ANN, word2vec, motion energy; **Figure 3**). ANN features showed the largest correlations in the bilateral IPS, in line with previous studies on math cognition (Nieder 2016; Arsalidou and Taylor 2011). In contrast to ANN features, word2vec (i.e. language) features showed the largest correlation in the left precentral sulcus (slightly posterior to the inferior frontal gyrus) and left-lateralised correlation patterns, whereas motion energy (i.e. visual) features showed the largest correlation in the bilateral occipital cortex. These results are consistent with previous reports on language and visual neurosciences (Bradshaw et al. 2017; Nishimoto et al. 2011; Güçlü and van Gerven 2015), suggesting that different latent feature distributions represent distinct aspects of mathematical operations.

The FBS analysis enabled the reconstruction of a brain representation of the sparse operator model based on the latent ANN model (**Figure 4**). Interestingly, the ANN model outperformed the other two latent feature models across whole-cortical voxels. Although it might seem contradictory to the result with RSA, difference between analyses might be due to different analytic procedures. The RSA calculates correlations across different operators for each anatomical ROI, whereas the FBS calculates correlations separately for each operator in each cortical voxel. Therefore, the FBS can reveal the finer organisation of the distributed information of mathematical operations in the brain, and is a promising way to bridge the gap between neuroimaging based on sparse coding and population coding (Foldiak 2003; Thorpe 2012).

The increase in correlation between operator-weight maps and FBS maps may reflect a representational hierarchy of mathematical operations in the ANN model (**Figure 4F, S4**). Previous studies revealed a correspondence of representational hierarchy between ANNs and BNNs in the visual (Yamins et al. 2014; Güçlü and van Gerven 2015; Horikawa and Kamitani 2017), auditory (Kell et al. 2018; Koumura, Terashima, and Furukawa 2019) and language domains (Hannagan et al. 2021; Caucheteux and King 2022). Our result is in line with these previous works, in that deeper ANN layers represent latent features underlying abstract cognitive processes such as mathematical operations.

In contrast to previous studies on operator decoding (Pinheiro-Chagas, Piazza, and Dehaene 2019; Kutter et al. 2022; Knops et al. 2009; Haynes et al. 2007), we successfully decoded novel operators (**Figure 5**). Such generalisation was achieved by using latent ANN features. Our models also showed generalised decodability across math expressions with different structures (i.e. single- and double-operator problems). Additional analyses using decoding models trained without target operators demonstrated that the decoding results were not a product of shared operators in math expressions (e.g. ‘AddMul’ and ‘AddDiv’ share ‘Add’). These results indicate that the ANN model represents the brain activity pattern of mathematical operations as distributions in a high-dimensional latent feature space.

It is worth noting some limitations in the current study. First, we did not compare the performance of different ANNs for math problem solving (e.g. (Wang et al. 2019; Andor et al. 2019; Lample and Charton 2019). However, the current study did not aim to determine the best fitting ANN model for the brain and TP-transformer is advantageous as an interpretation of its latent vectors has already been reported (Russin et al. 2021). The best brain-like ANN model of mathematical operations should be determined in future studies using concepts such as the brain-score (Schrimpf et al. 2020). Second, although the number of subjects was determined based on previous studies using encoding/decoding (Huth et al. 2016; Nakai and Nishimoto 2020; Popham et al. 2021; Horikawa et al. 2020), the relatively small number of subjects involved may affect the current findings. Indeed, two subjects out of eight showed inconsistent patterns of prediction map similarity (**Figure 2C, D**). These subjects, however, also showed a significant correlation between FBS and operator-weight maps (**Figure 4E**), indicating that the representational relationship between the sparse operator and latent ANN models is robust across individuals. Further research on a larger dataset may clarify individual differences in the distributed brain representations of mathematical operations. The current study is a first step toward ANN modelling of the brain representations of complex human cognitive abilities, such as mathematical thought, bridging the gap between symbolic and distributed representations found in human neuroimaging.

## Methods

### Subjects

Eight healthy college students (aged 20–23 years, three females, all with normal vision), denoted as ID01–ID08, participated in this study. This number is in the same range as previous encoding model studies (Huth et al. 2016; Nakai and Nishimoto 2020; Popham et al. 2021; Horikawa et al. 2020). Subjects were all right-handed (laterality quotient, 80–100) as assessed using the Edinburgh inventory (Oldfield 1971). Written informed consent was obtained from all subjects prior to their participation in the study; the study was approved by the ethics and safety committee of the National Institute of Information and Communications Technology in Osaka, Japan.

### Stimuli and procedure

We selected arithmetic problems with a single operation, addition (Add), subtraction (Sub), multiplication (Mul) and division (Div), and arithmetic problems with two operations, including addition and subtraction (AddSub), addition and multiplication (AddMul), addition and division (AddDiv), subtraction and multiplication (SubMul) and subtraction and division (SubDiv). Each condition consisted of 35 instances. Subjects also underwent another math problem-solving task in word format and a control sentence comprehension task; these tasks were analysed elsewhere (Nakai and Nishimoto 2022). Behavioural data were also analysed in this previous work.

In each trial, an arithmetic expression problem (e.g. ‘3 × 2 =?’) was presented for 4 or 6 s (4 s for single-operator conditions, and 6 s for double-operator conditions). The subjects were instructed to perform a calculation based on the presented problem and to press the button on the left when they had an answer. The fixation cross stimulus was then presented for 1–2 s, after which a probe digit stimulus was presented for 2 s (e.g. ‘6’). Subjects were again asked to press the left or right button if the presented digit matched or did not match their answer, respectively. The next trial began after the fixation cross was again displayed for 1–2 s. The number of letters in each condition was as follows: 5.5 ± 0.5 (Add), 6.5 ± 0.5 (Sub), 5.0 ± 0.2 (Mul), 5.9 ± 0.4 (Div), 9.3 ± 0.9 (AddSub), 8.9 ± 0.3 (AddMul), 10.1 ± 0.7 (AddDiv), 9.6 ± 0.5 (SubMul) and 10.9 ± 0.5 (SubDiv).

Stimuli were presented on a projector screen inside the scanner (21.0 × 15.8° visual angle at 30 Hz). During scanning, subjects wore MR-compatible ear tips. The experiment was performed over three days, with 5–7 runs per each day. A presentation software (Neurobehavioral Systems, Albany, CA, USA) was used to control stimulus presentation and collection of behavioural data. Optic response pads with two buttons were used to measure button responses (HHSC-2×2, Current Designs, Philadelphia, PA, USA).

Six training and two test runs were conducted. The data from each pair of test runs were averaged to increase the signal-to-noise ratio. Each run contained 90 trials. A single run lasted 455 s. At the beginning of each run, we acquired 10 s of dummy scans, during which the fixation cross was displayed; these dummy scans were later omitted from the final analysis to reduce noise. We also obtained 10 s of scans at the end of each run, during which the fixation cross was displayed, and which were included in the analyses.

### MRI data acquisition

The experiment was conducted using a 3.0 T scanner (MAGNETOM Prisma; Siemens, Erlangen, Germany) with a 64-channel head coil. We scanned 72 interleaved axial slices, 2-mm thick without a gap, parallel to the anterior and posterior commissure line, using a T2*-weighted gradient-echo multiband echo-planar imaging sequence [repetition time (TR) = 1,000 ms; echo time (TE) = 30 ms; flip angle (FA) = 62°; field of view (FOV) = 192 × 192 mm^2^; resolution = 2 × 2 mm^2^; multiband factor = 6]. We obtained 605 volumes for the MW and RW conditions and 465 volumes for the ME condition, each set following 10 dummy images. As anatomical reference, high-resolution T1-weighted images of the whole brain were acquired from all subjects with a magnetisation-prepared rapid acquisition gradient-echo sequence (TR = 2,530 ms; TE = 3.26 ms; FA = 9°; FOV = 256 × 256 mm^2^; voxel size = 1 × 1 × 1 mm^3^).

### fMRI data pre-processing

In each run, motion correction was performed using the statistical parametric mapping toolbox (SPM12; Wellcome Trust Centre for Neuroimaging, London, UK; http://www.fil.ion.ucl.ac.uk/spm/). All volumes were aligned to the first EPI image for each subject. Low-frequency drift was removed using a median filter with a 120-s window. Slice timing correction was performed against the first slice of each scan. The response for each voxel was then normalised by subtracting the mean response and scaling it to the unit variance. We used FreeSurfer (https://surfer.nmr.mgh.harvard.edu/) to identify the cortical surfaces from anatomical data and register these to functional data voxels. For each subject, identified voxels in the cerebral cortex (59,499–75,980 voxels per subject) were used for analysis.

### Operator features

Operator features included one-hot vectors with assigned values of 1 or 0 for each time bin during the stimulus presentation of arithmetic problems, indicating whether one of the nine tested conditions (Add, Sub, Mul, Div, AddSub, AddMul, AddDiv, SubMul and SubDiv) was performed in that period; thus, nine operator features were used.

### ANN features

ANN features were extracted using a pretrained TP-transformer model (Schlag et al. 2019). TP-Transformer, an extension of the transformer model (Vaswani et al. 2017) consisting of encoder and decoder layers, where the former layer processes given math expressions and the latter generates the answer. Both encoder and decoder have six transformer layers containing multi-head attention modules with eight heads. In contrast to the regular transformer where the attention head consists of query, key and value vectors, TP-Transformer additionally uses a role vector to better capture structural information in the input math expressions. Each attention head thus consists of query (Q), key (K), value (V) and roll (R) vectors for each input symbol:

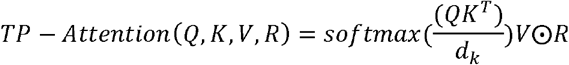

where dk is a dimension of K vector and ⊙ is Hadamard product. The other part of the architecture and hyper-parameters of the TP-Transformer is similar to the regular transformer (dimensionality of input and output, *d_model_* = 512; dimensionality of feed-forward network, *d_f_* = 2048). The model was trained using the Mathematics Dataset (Saxton et al. 2019).

For each input math expression (e.g. ‘3 + 5’), we extracted the average query, key, value and roll vectors from each of the six encoding layers. Time points without vector assignments were defined as 0. Although we could use all these vectors, we used the query vectors of the sixth encoding layer as a representative vector based on a previous study (Russin et al. 2021). We also extracted query vectors of the sixth encoding layer with each mathematical symbol (e.g. ‘3’, ‘+’, etc.) as input. ANN features had 512 dimensions.

### ME features

We employed a previously described ME model (Nishimoto et al. 2011; Nakai and Nishimoto 2020; Koide-Majima, Nakai, and Nishimoto 2020) available in a public repository (https://github.com/gallantlab/motion_energy_matlab). First, movie frames and pictures were spatially downsampled to 96 × 96 pixels. RGB pixel values were then converted into the Commission International de l’Eclairage (CIE) LAB colour space; then, colour information was discarded. The luminance (L*) pattern was passed through a bank of three-dimensional spatiotemporal Gabor wavelet filters; the outputs of two filters with orthogonal phases (quadrature pairs) were squared and summed to yield the local ME. Subsequently, MoE was compressed with a log-transformation and temporally downsampled to 0.5 Hz. Filters were tuned to six spatial (0, 1.5, 3.0, 6.0, 12.0 and 24.0 cycles per image) and three temporal frequencies (0, 4.0 and 8.0 Hz) without directional parameters. Filters were positioned on a square grid covering the screen. Adjacent filters were separated by 3.5 standard deviations of their spatial Gaussian envelopes. To reduce the computational load, the original ME features, which had 1,395 dimensions, were reduced to 300 dimensions using principal component analysis.

### Word2vec features

To quantitatively evaluate the brain representations of the presented semantic information in a data-driven manner, we extracted the semantic features from each narrative stimulus using word2vec (Mikolov et al. 2013). Word2vec, based on the skip-gram model, has been used to embed words into distributed representations. All math expressions were segmented into words (i.e. symbols) and morphologically parsed using MeCab (https://taku910.github.io/mecab/). Individual word segments were projected into the 300-dimensional space. Timepoints without any word vector assignments were defined as 0. The resultant concatenated vectors were downsampled to 1 Hz. The dimension size of word vectors was set to the default value of 300.

### BR feature

The single button response (BR) feature was constructed based on the number of button responses per second.

### Letter feature

The single letter feature was constructed based on the number of letters appearing in each stimulus.

### RT feature

For each time bin during presentation of mathematical problems, the single trial RT was assigned as the trial’s cognitive load.

### Encoding model fitting

In the encoding model, the cortical activity in each voxel was fitted with a finite impulse response model that captured the slow hemodynamic response and its coupling with neural activity (Nishimoto et al. 2011; Kay, David, et al. 2008). The feature matrix **F_E_** [T × 6N] was modelled by concatenating sets of [T × N] feature matrices with six temporal delays between 2–7 s (T = number of samples; N = number of features). The cortical response **R_E_** [T × V] was then modelled by multiplying the feature matrix **F_E_** by the weight matrix **W_E_** [6N × V] (V = number of voxels):

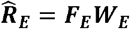

We used L2-regularised linear regression with the training dataset (consisting of 2,790 [2,790 s] samples) to obtain the weight matrix **W_E_**. The optimal regularisation parameter was assessed using 10-fold cross-validation where the 11 different regularisation parameters ranged from 1 to 2^10^.

The test dataset consisted of 465 samples (465 s, repeated twice). Two repetitions of the test dataset were averaged to increase the signal-to-noise ratio. Statistical significance (one-sided) was computed by comparing estimated correlations to the null distribution of correlations between two independent Gaussian random vectors with the same length as the test dataset. The statistical threshold was set at *P* < 0.05 and corrected for multiple comparisons using FDR (Benjamini and Hochberg 1995). For data visualisation on cortical maps, we used pycortex (Gao et al. 2015) and fsbrain (Schäfer and Ecker 2020).

### Encoding model fitting excluding regressors of noninterest

To exclude the possible effect of sensorimotor processing, linguistic processing and general cognitive load on the model predictions, we concatenated ME (visual), word2vec (semantic), BR; motor), RT (cognitive load) and letter (orthographic) features to the original operator and ANN feature matrices. The concatenated features were used as a new feature matrix for modelling encoding. L2-regularised linear regressions were applied as described in the ‘Encoding model fitting’ subsection. For model testing, prediction accuracy was evaluated using the feature matrix after excluding non-interesting features.

### RSA

To compare ANN features with the sparse operator model, we performed RSA based on two different RSMs. First, we calculated the ANN-to-Operator transform matrix (AOTM) using the average features extracted from TP-Transformer for all trials in the training dataset. A feature-brain RSM was calculated using the Pearson correlation distance across all combinations of the nine operators in the AOTM. Second, for each operator, we averaged the weight vectors of the operator encoding model for all voxels included in each target anatomical ROI; six-time delays were further averaged. A brain-based RSM was calculated using the Pearson correlation distance across all combinations of nine operators in the resultant mean weight vectors. The upper triangular parts of the feature-based and brain-based RSMs were rearranged into single vectors and the Spearman’s correlation coefficients between the two calculated.

### FBS analysis

To provide voxel-wise similarity information and differences in patterns across operators, we calculated the FBS for each cortical voxel by calculating the Pearson’s correlation coefficient between the reference latent feature vector and the voxel-specific weight vector in each cortical voxel. The reference latent feature vector was taken from the AOTM calculated similarly to the RSA. The voxel-specific weight vector was taken from each target model weight matrix and averaged for six temporal delays.

### Decoding model fitting

In the decoding model, the cortical response matrix R_D_ [T × 6V] was modelled using concatenating sets of [T × V] matrices with temporal delays of 2–7 s. The feature matrix F_D_ [T × N] was modelled by multiplying the cortical response matrix R_D_ by the weight matrix W_D_ [6V × N]:

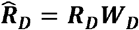

The weight matrix WD was estimated using an L2-regularised linear regression with the training dataset, following the same procedure as for encoding model fitting.

To examine the generalizability of our models, we performed three types of novel operator decoding. (1) First, we performed nine independent model fittings, each with a different operator as target. From the training dataset, we excluded the time points during which the target condition (i.e. problems with the target operator) occurred and those within 7 s after presentation of the target condition. In the test dataset, we used only the time points during which the target condition occurred and those within 7 s after presentation of the target condition. This allowed us to assume that the activity induced by the target condition and that induced by the other eight operator groups (training conditions) did not overlap, and to investigate prediction and decoding accuracies for novel operators. (2) Second, we performed two independent model fittings, where only single- or double-operator problems were used as training dataset; the remaining operators (double-operators in case of single operators used in model training) were used in the model testing phase. (3) Third, we performed four independent model fittings for four single operators (‘Add’, ‘Sub’, ‘Mul’ and ‘Div’) as target, where problems without target operator (e.g. without ‘Add’, ‘AddSub’, ‘AddMul’ and ‘AddDiv’ in the case of ‘Add’ as a target); the target operator (e.g. ‘Add’) was used in the model testing phase.

For the evaluation of decoding accuracy, we measured the similarity between the AOTMs of each condition and each decoded vector using the Pearson’s correlation coefficients for each time point. We refer to the correlation coefficient as *task score* (Nishida and Nishimoto 2018). We then plotted a ROC curve for each target operator by plotting the ratio of true and false positives by changing the discrimination threshold based on the task score. Decoding accuracy was evaluated using the AUC. The statistical significance of the decoding accuracy for each condition was tested using a one-sided permutation test *(p* < 0.05, with FDR correction).

## Data and code availability

The source data and analysis code used in the current study are available from Zenodo (https://doi.org/XXXXX).

## Acknowledgements

We thank MEXT/JSPS KAKENHI (grant numbers JP17K13083 and JP18H05091 in #4903 (Evolinguistics) for T.N., JP15H05311 and JP18H05522 for S.N.) as well as JST CREST JPMJCR18A5 and ERATO JPMJER1801 (for S.N.) for the partial financial support of this study. The funders had no role in the study design, data collection and analysis, decision to publish, or preparation of the manuscript.

## Author contributions

T.N. designed the study; T.N. collected and analysed the data; T.N. and S.N. wrote the manuscript.

## Supplementary Information

**Figure S1.**
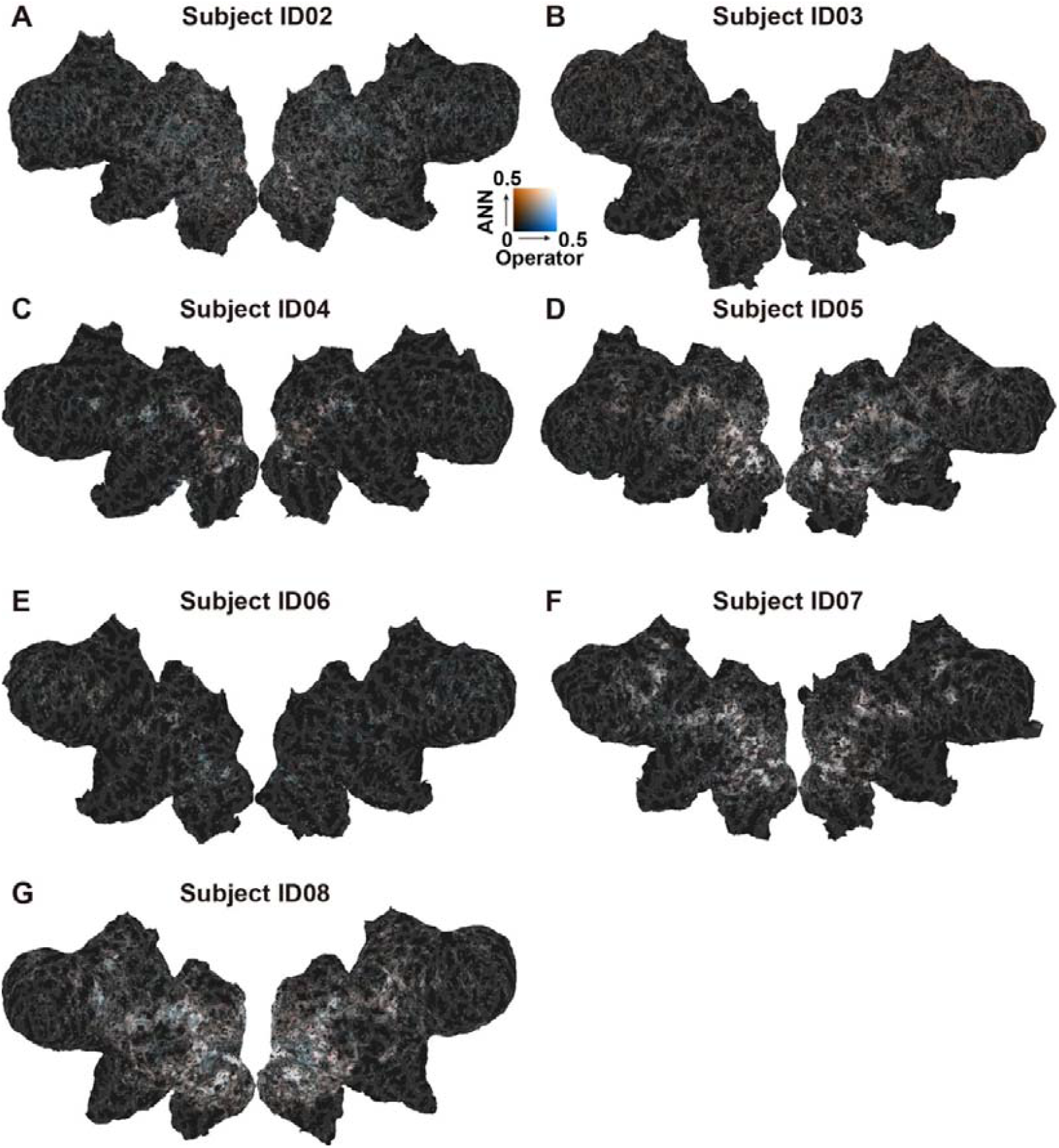
Prediction of brain activity by ANN and operator encoding models. Prediction accuracies of ANN (red) and operator (blue) models shown on the cortical surface of subject ID02-ID08. Only voxels with significant prediction are shown *(p* < 0.05, FDR corrected).

**Figure S2.**
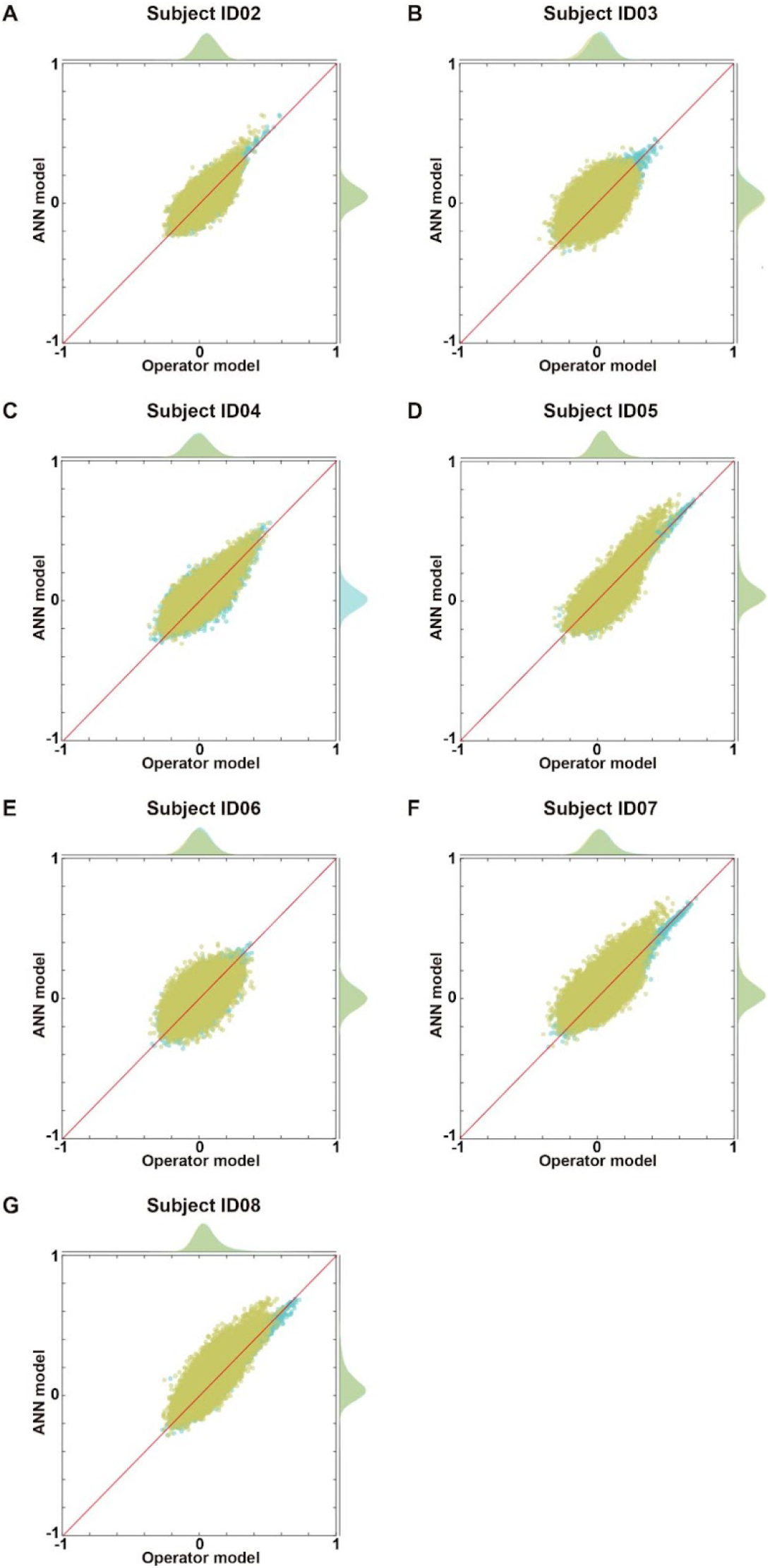
Prediction of brain activity by ANN and operator encoding models. Scatter plot of ANN and operator models for subject ID02-ID08, shown for both original model (cyan) and after exclusion of non-mathematical regressors (yellow).

**Figure S3.**
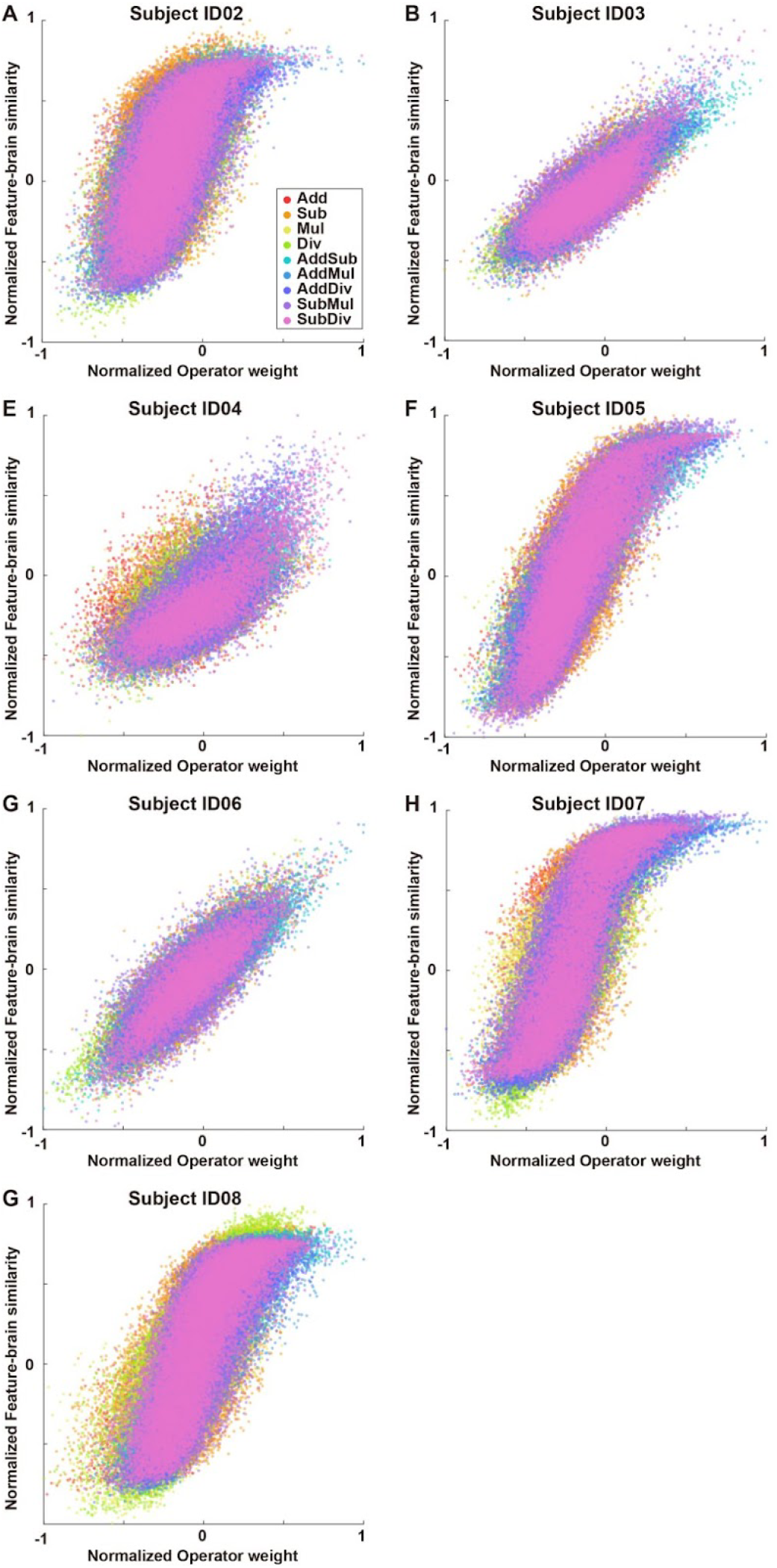
Correlation between operator weight and FBS values. Scatter plot of all cortical voxels showing a correlation between operator weight and FBS values for subject ID02-ID08.

**Figure S4.**
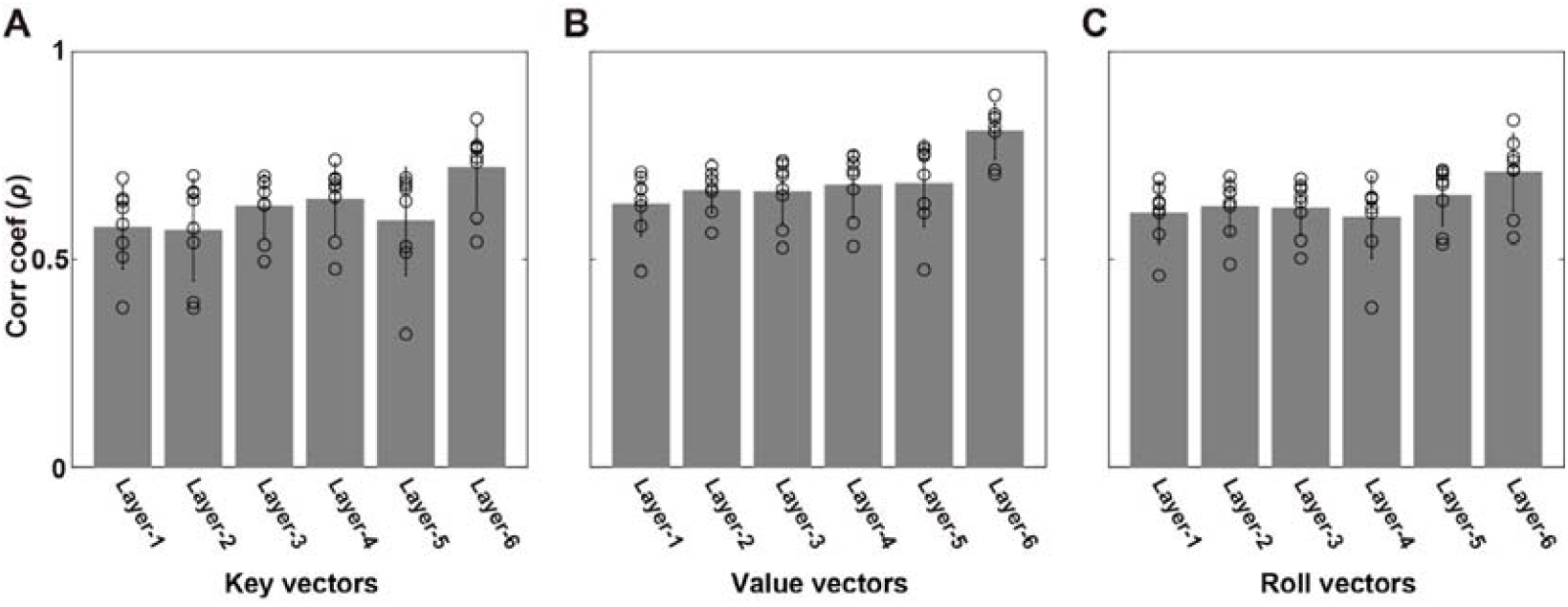
Correlations of operator-weight maps and FBS maps for different ANN layers. Mean Spearman’s correlation coefficients between operator weights and FBS values were obtained using (A) key, (B) value and (C) roll vectors from different ANN encoding layers (layers 1 to 6). Individual subjects’ data are indicated by small circles. Error bar, SD.

**Figure S5.**
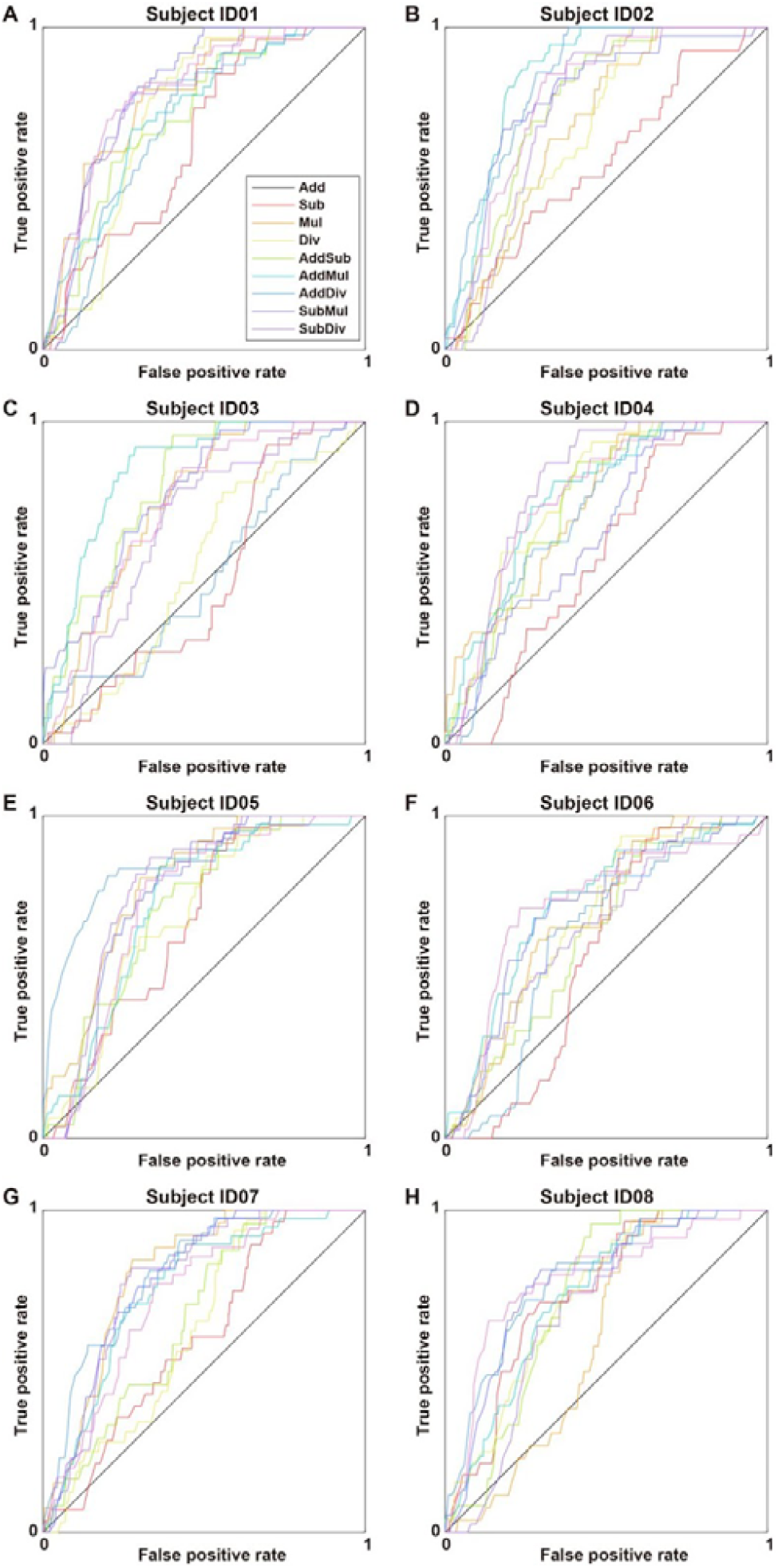
Receiver operating characteristic curves. Receiver operating characteristic curves were plotted for subject ID02-ID08 based on ANN decoding models trained with eight of nine operators and tested with the remaining novel operator.

